# The core genome ^m5^C methyltransferase JHP1050 (M.Hpy99III) plays an important role in orchestrating gene expression in *Helicobacter pylori*

**DOI:** 10.1101/393710

**Authors:** Iratxe Estibariz, Annemarie Overmann, Florent Ailloud, Juliane Krebes, Josenhans Josenhans, Sebastian Suerbaum

**Author notes:** To whom correspondence should be addressed.; Fax: -4989218072802; Email: Christine Josenhans or Sebastian Suerbaum.

## Abstract

*Helicobacter pylori* encodes a large number of Restriction-Modification (R-M) systems despite its small genome.R-M systems have been described as “primitive immune systems” in bacteria, but the role of methylation in bacterial gene regulation and other processes is increasingly accepted. Every *H.pylori* strain harbours a unique set of R-M systems resulting in a highly diverse methylome. We identified a highly conserved GCGC-specific ^m5^C MTase (JHP1050) that was predicted to be active in all of 459 *H.pylori* genome sequences analyzed. Transcriptome analysis of two *H.pylori* strains and their respective MTase mutants showed that inactivation of the MTase led to changes in the expression of 225 genes in strain J99, and 29 genes in strain BCM-300.10 genes were differentially expressed in both mutated strains. Combining bioinformatic analysis and site-directed mutagenesis, we demonstrated that motifs overlapping the promoter influence the expression of genes directly, while methylation of other motifs might cause secondary effects.Thus, ^m5^C methylation modifies the transcription of multiple genes, affecting important phenotypic traits that include adherence to host cells, natural competence for DNA uptake, bacterial cell shape, and susceptibility to copper.

## INTRODUCTION

Epigenetics denotes inheritable mechanisms that regulate gene expression without altering the DNA sequence. In prokaryotes, methyltransferases (MTases) transfer methyl groups from S-adenosyl methionine to adenines or cytosines within a DNA target motif and so contribute to changes of the epigenome (1-3). MTases either belong to Restriction-Modification (R-M) systems that include MTase and restriction endonuclease (REase) activities, or occur as orphan MTases in the absence of a cognate restriction enzyme (4). Three types of DNA methylation occur in bacteria, N6-methyladenine (^m6^A), 5-methylcytosine (^m5^C) and N4-methylcytosine (^m4^C) (1,2). So far, the major role allocated to bacterial R-M systems is self-DNA protection by restriction of incoming foreign un-methylated DNA (5), and they have thus been described as “primitive immune systems” (6). Other functions have also been attributed to prokaryotic R-M systems (7-9). For example, methylation marks promoter sequences and alters DNA stability and structure, modifying the affinity of DNA binding proteins and influencing the expression of genes (10,11). Additionally, disturbance of DNA strand separation by methylation can have an effect on gene expression (12).

Methylation can be involved in multiple bacterial functions. In *Escherichia coli*, the Dam adenine MTase plays an essential role in DNA replication (13,14). Another well-studied example is the CcrM MTase from *Caulobacter crescentus* that controls the progression of the cell cycle (15). Furthermore, phase-variable MTases have been shown to control the regulation of multiple genes in several different pathogens, including *Haemophilus influenzae, Neisseria meningitidis*, and *Helicobacter pylori* (16-18). These MTase-dependent regulons were termed phasevarions (19). As described previously, adenine methylation has been shown to play a key role in transcriptional regulation but the influence of cytosine methylation in gene expression has so far only been investigated in very few studies (20,21).

*H. pylori* infection affects half of the world’s population and is a major cause of gastric diseases that include ulcers, gastric cancer, and MALT lymphoma (22). This gastric pathogen has coexisted with humans since, at least, 88,000 years ago (23).*H. pylori* strains display an extraordinary genetic diversity caused in part by a high mutation rate but especially by DNA recombination occurring during mixed infection with other *H. pylori* strains within the same stomach (24-26). The very high sequence diversity of *H. pylori* and the coevolution of this pathogen with its human host have caused its separation into phylogeographic populations, whose distribution reflects human migrations (27-29).

Despite its small genome, *H. pylori* is one of the pathogens with the highest number of R-M systems (30). The development of Single Molecule, Real-Time (SMRT) Sequencing technology has allowed genome-wide studies of methylation patterns and strongly accelerated the functional elucidation of MTases and their roles in bacterial biology (31,32). Methylome studies of several *H. pylori* strains have revealed that every strain carries a different set of R-M systems leading to highly diverse methylomes (33-36). R-M systems in *H. pylori* were shown to protect the bacterial chromosome against the integration of non-homologous DNA (e. g. antibiotic resistance cassettes), while they had no significant effect on recombination between highly homologous sequences, permitting efficient allelic replacement (9). Despite the diversity of methylation patterns, a small number of target motifs were shown to be methylated in all (one motif, GCGC) or almost all (3 motifs protected in >99% of strains) *H. pylori* strains in a study by Vale *et al*., who tested genomic DNAs purified from 221 *H. pylori* strains for susceptibility to cleavage by 29 methylation-sensitive restriction enzymes, and in those studies investigating the methylomes of multiple *H. pylori* strains (33,34,36,37). R-M systems have also previously been shown to contribute to gene regulation in *H. pylori;* the phase-variable MTase ModH5 is involved in the control of the expression of virulence-associated genes like *hopG* or *flaA* in strain P12 (38,39).

In the present study, we functionally characterized the role of a highly conserved ^m5^C MTase (JHP1050, M. Hpy99III) in *H. pylori* (40). We show the MTase gene to be part of the *H. pylori* core genome, present and predicted to be active in all of several hundred *H. pylori* strains representative of all known phylogeographic populations. Transcriptome comparisons of two *H. pylori* wild-type strains and their respective knockout mutants demonstrated that JHP1050 has a strong impact on the *H. pylori* transcriptome that includes both conserved and strain-specific regulatory effects. We show that methylation of G^m5^CGC sequences, among others, affects metabolic pathways, competence and adherence to gastric epithelial cells. Moreover, we provide specific evidence that methylation of motifs within promoter sequences can play a direct role in gene expression, while the regulatory effects of methylated sites outside of promoter region may be indirect.

## MATERIAL AND METHODS

### Bacterial culture, growth curves and transformation experiments

*H. pylori* strains were cultured on blood agar plates (41), or in liquid cultures as described (9). Microaerobic conditions were generated in airtight jars (Oxoid, Wesel, Germany) with Anaerocult C gas producing bags (Merck, Darmstadt, Germany). For growth curves, liquid cultures were inoculated with bacteria grown on agar plates for 22-24 h to a starting OD_600_ of ~0.06 and incubated with shaking (37°C, 140 rpm, microaerobic conditions). The OD_600_ was repeatedly measured until a maximum incubation time of 72 hours.

Susceptibility to copper was tested by adding copper sulphate (final concentrations, 0.25 mM and 0.50 mM) to liquid cultures. The OD_600_ was measured 24 hours after inoculation. For transformation experiments, liquid cultures of the recipient strain were grown overnight (conditions described above). Then, 1 μg/ml of donor bacterial genomic DNA (gDNA) was added to the cultures. The donor gDNA for transformation experiments was purified from isogenic *H. pylori* strains carrying a chloramphenicol (CAT) resistance cassette within the non-essential *rdxA* gene (i. e. J99 *rdxA∷*CAT). After gDNA addition, the cultures were incubated for 6-8 hours under the same conditions (37°C, 140 rpm, microaerobic atmosphere). Next, the OD_600_ was measured and adjusted to the same number of cells (OD_600_ = 1 as 3 × 10^8^ bacteria). Finally, 100 μl of serial dilutions were plated onto blood agar plates containing chloramphenicol, and incubated at 37°C under microaerobic conditions. Approximately 4-5 days later, colonies were counted and the efficiency of transformation was calculated as cfu/ml.

### DNA and RNA extraction

gDNA was isolated from bacteria grown on blood agar plates using the Genomic-tip 100/G kit (Qiagen, Hilden, Germany) following the manufacturer’s protocol. The gDNA pellet was dissolved over night at room temperature with EB buffer.

For RNA extraction, 5 ml of bacterial cells grown in liquid medium were pelleted (4°C, 6000 x g, 3 min), snap-frozen in liquid nitrogen and stored at -80 °C. Afterwards, bacterial pellets were disrupted with a FastPrep®FP120 Cell Disrupter (Thermo Savant) using Lysing Matrix B 2 ml tubes containing 0.1 mm silica beads (MP Biomedicals, Eschwege, Germany). Isolation of RNA was performed using the RNeasy kit (Qiagen, Hilden, Germany) and on-column DNase digestion with DNase I. A second DNase treatment was carried out using the TURBO DNA-free^TM^ Kit (Ambion, Kaufungen, Germany). Isolated RNA was checked for the absence of DNA contamination by PCR reaction.

DNA and RNA concentrations were measured using a NanoDrop 2000 spectrophotometer (Peqlab Biotechnologies). RNA quality given as RINe number was measured with an Agilent 4200 Tape Station system using RNA Screen Tapes (Agilent, Waldbronn, Germany). All the RINe numbers were higher than 8.2, suggesting a low amount of degradation products.

### Construction of mutants and complementation

I nactivation of the MT ase or the whole R-M system genes was carried out by insertion of an *aphA3* cassette conferring resistance to kanamycin (Km). A PCR product was constructed using a combination of primers which added restriction sites and allowed overlap PCR with the *aphA3* cassette (Q5 Polymerase, NEB, Frankfurt am Main, Germany). Ligation of the overlap amplicon with a digested pUC19 vector was done using the Quick Ligase (NEB, Frankfurt am Main, Germany). The resulting plasmids were transformed into *E. coli* MC1061. Following plasmid isolation, 750 ng of the plasmids were used for *H. pylori* transformation. Functional complementation of the MTase gene in the strains 26695-mut and J99-mut was achieved by means of the pADC/CAT suicide plasmid approach, as described (42). Transformation of the recipient strains with the resulting plasmids permitted the chromosomal integration of the MTase gene (from 26695) into the urease locus, placing the inserted gene under the control of the strong promoter of the *H. pylori* urease operon. The complemented strains are designated 26695-compl and J99-compl, respectively.

Domain mutants carrying point mutations within GCGC motifs were constructed using the Multiplex Genome editing (MuGent) technique as described (9,43), with the exception that we used only a CAT cassette within the non-essential *rdxA* locus as selective marker. Sanger sequencing was used to verify the acquisition of the desired mutations within the GCGC motifs. The putative promoter of the gene was predicted using the BPROM Softberry online tool (44). All *H. pylori* mutants were checked via PCR and selected on antibiotic-containing plates. The absence or recovery of methylation was checked by digestion of gDNA with HhaI (NEB, Frankfurt am Main, Germany). All plasmids and primers used in this study are listed on the Supplementary Tables 6 and 7.

### Microscopy

Live and Dead (L/D) staining was performed using the BacLight Bacterial Viability kit (Thermo Fisher Scientific, Darmstadt, Germany) according to the manufacturer’s instructions. Bacteria were harvested from plates incubated for 22-24 h, and suspended in 1 ml of BHI medium without serum to an adjusted OD_600_ of ~0.1. Then, 100 μl of this dilution were mixed with the BacLight dyes, giving green and red fluorescence for live and dead/dying bacteria, respectively. After 30 minutes of incubation at room temperature and in the dark, 0.5 μl of the mix was suspended on slides that were analyzed with an Olympus BX61-UCB microscope equipped with an Olympus DP74 digital camera. Between 80 and 100 pictures from at least two independent biological and technical replicates were obtained and analyzed with the CellSens 1.17 software (Olympus Life Science) and ImageJ (45).

Gram staining was performed as follows: 300 μl of liquid cultures grown over-night were pelleted (6,000 x g, 3 minutes, room temperature) and washed 3 times with PBS (6,000 x g, 3 minutes, room temperature). Afterwards, 100 μl of the pellets suspended in PBS were added to a glass slide that was dried at 37°C during 10-15 minutes, heat-fixed, and Gram-stained.

### Bacterial cell adherence assays

Cell adherence assays were performed as previously described with slight modifications (46,47).*H. pylori* strains grown to an OD_600_ ~1 were suspended in RPMI 1640 medium supplemented with 10% fetal calf serum (FCS). Experiments were executed in 96 well plates containing 2×10^5^ fixed AGS cells (ATCC CRL-1739) per well. AGS cells were fixed with 2% paraformaldehyde in 100 mM potassium phosphate buffer (pH=7) and subsequently quenched and washed as described (46). Live *H. pylori* bacteria were added to cells at a bacteria:cell ratio of 50 (47), followed by brief centrifugation (300 x g, 5 min), and co-incubated for 1 h at 37°C with 5% CO_2_. After this, plates were washed twice with PBS, followed by overnight fixation with 50 μl of fixing solution (see above). Fixing solution was renewed once and incubated for an additional 30 min, and quenched twice with 50 μl of quenching buffer for 15 min. Bacterial adherence to the AGS cells was quantitated as follows: cells were washed three times with washing buffer PBS-T (PBS + 0.05% Tween20), blocked for 30 min with 200 μl of the assay diluent (10% FCS in PBS-T) and washed four times with PBS-T. Then, 100 μl of a 1:2,500 dilution of the primary antibody α-H. pylori (DAKO/Agilent, Hamburg, Germany) were added and incubated for 2 h. Afterwards, cells were washed and incubated with 100 μl of a 1:10,000 dilution of the secondary antibody, goat anti-rabbit HRP (Jackson ImmunoResearch, Ely, United Kingdom) for 1 h. After four final wash steps, the 96-well plates were finally incubated with 100 μl TMB substrate solution (1:1, Thermo Fisher Scientific, Darmstadt, Germany). The reaction was developed in the dark for 30 min and stopped with 50 μl of phosphoric acid (1 M). Absorbance was measured at 450/540 nm (Sunrise^TM^ Absorbance Reader). Negative controls (mock-coincubated, fixed AGS cells) were treated the same way with primary and secondary antibody dilutions.

### Bioinformatic analyses

To analyze the conservation and the genomic context of the JHP1050 MTase gene in a diverse collection of *H. pylori* strains, we assembled a database consisting of 459 *H. pylori* genomes that included strains from all known phylogeographic populations and subpopulations (Supplementary Table 1). The nucleotide sequence of gene *jhp1050* from the *H. pylori* strain J99 was used to identify and extract the *jhp1050* homologs and the sequences of the flanking genes. The NCBI blastn microbes and StandAlone Blast tools were used to extract the sequences from publicly available genomes and private genomes, respectively.

T o study whether the methylated cytosines of the GCGC motifs had a higher tendency to deaminate (^m5^C>T) than unmethylated cytosines, we compared the frequency of C>T transitions to either C>A or C>G polymorphisms inside and outside of GCGC motifs among a phylogeographically distinct set of *H. pylori* genomes. GCGC motifs were identified in two *H. pylori* genomes, 26695 and PeCan18, which were subsequently used as reference and aligned separately against 11 other *H. pylori* genomes using BioNumerics version 7.6 (Applied Maths, Sint-Martens-Latem, Belgium). Polymorphisms were called in both alignments and pooled together. The percentage of mutated ^m5^C positions within GCGC motifs was determined for each possible transition or transversion as follows:

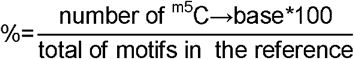

Since the G^m5^C<u>G</u>C motif is palindromic, the same analysis was made for the complementary strand, where the position of the second G (^m5^C in the complementary strand) was compared for each possible mutation and calculated as above.
Finally, the percentage of mutated C outside GCGC motifs calculated as follows for each possible mutation:

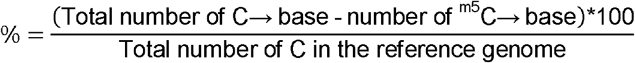

### Predicted sites

The predicted number of motifs per kb was calculated as follows:

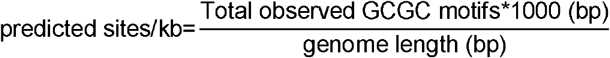

The predicted number of motifs/kb was 3.89 (J99), 3.91 (BCM-300), 3.76 (26695) and 3.74 (H1). The expected number of motifs within CDS can then be calculated using the predicted number of motifs/kb and the gene length. Finally, the ratio observed/expected (O/E) motifs within CDS can be calculated to detected genes enriched for motifs presence. For example; a given gene in J99 that is 630 bp long and has two GCGC motifs (observed). The expected number of motifs within that gene would be: 630*3.89/1000 (expected). The ratio O/E would then be 0.8161 suggesting GCGC motifs are underrepresented in this gene.

The GCGC motif is a 4-mer palindrome. In order to calculate the expected number of motifs that would randomly occur within the genome, the CDS and intergenic regions, we took into account the number of 4-mers in a given sequence, N-K+1 (where N means sequence length and K the motif length, in this case 4); and the frequency of G/C (0.2) and A/T (0.3). The final formula would be: (N-K+1)*(0.2)^4^.

### RNA-Seq analysis

RNA-Seq analysis was performed on an Illumina HiSeq sequencer obtaining single end reads of 50 bp. Total rRNA depletion was performed prior to cDNA synthesis using a RiboZero Kit (Illumina, Germany). Isolated RNA from a total of 6×10^8^ – 1×10^9^ bacterial cells corresponding to log phase of growth was used for sequencing. Three biological replicates were used for all the strains, except for J99-mut since one replicate had to be discarded during library preparation. Mapping of reads to a reference genome was done with Geneious 11.0.2 (48). Reads mapping multiple locations or intersecting multiple CDS were counted as partial matches (i. e.0.5 read). Differential expression was calculated using DESeq2 (49). Fold Change (FC) of two and FDR adjusted p-value of 0.01 were used as a cut-off.

RNA-seq data was placed in the ArrayExpress database at EMBL-EBI (www.ebi.ac.uk/arrayexpress) with accession number E-MTAB-xxxx.

### Quantitative PCR (qPCR)

One μg of RNA was used for cDNA synthesis using the SuperScript^TM^ II Reverse Transcriptase (Thermo Fisher Scientific, Darmstadt, Germany) as described before (47). qPCR was performed with gene specific primers (Supplementary Table 7) and SYBR Green Master Mix (Qiagen, Hilden, Germany). Reactions were run in a BioRad CFX96 system. Standard curves were produced and samples were run as technical triplicates. For quantitative comparisons, samples were normalized to an internal 16S rRNA control qPCR.

## RESULTS

### Distribution of the G^m5^CGC R-M system (JHP1049-1050) within a globally representative collection of *H. pylori* genomes

Despite the extensive inter-strain methylome diversity of *H. pylori*, a small number of motifs have been shown to be methylated in all or most of the strains (37). Here, we focused on the MTase JHP1050 (M. Hpy99III), which methylates GCGC sequences, resulting in G^m5^CGC motifs. Although ^m5^C methylation is less common in prokaryotes than ^m6^A-methylation, based on the Restriction Enzyme Database (REBASE) (50), this particular motif is highly conserved in many bacterial species. We therefore hypothesized that the Gm5CGC-specific MTase in *H. pylori* might play an important role apart from self-DNA protection.

We first analyzed the conservation and the genomic context of the MTase gene. The nucleotide sequence of gene *jhp1050* from the *H. pylori* strain J99 was used to identify the *jhp1050* homologs and the sequences of the flanking genes in a collection of 458 *H. pylori* genomes representing all known phylogeographic populations (Supplementary Table 1).

Based on the gene sequences, the M. Hpy99III MTase was predicted to be active in all *H. pylori* strains. The analyzed region of the chromosome was highly conserved among the strains and all the flanking genes were present with the exception of the cognate REase gene *(jhp1049*) which was present in only 61 of the 459 strains. Interestingly, the majority of the REase-positive strains belong to populations with substantial African ancestry, particularly to hpAfrica2, followed by hspSAfrica, hspWAfrica and hpEurope. Furthermore, none of the analyzed hspAsia2 or hspEAsia strains carried the REase gene (Supplementary Table 1). Only 15 REase genes were predicted to be functional, while the others were pseudogenes due to premature stop codons and/or frameshift mutations (Supplementary Table 2). We identified a 10 bp repeat sequence flanking the REase gene. The same sequence was found downstream of the MTase gene and 48 bp upstream of *jhp1048* in 15 of the REase-negative strains. In all cases, the sequence contained a homopolymeric region with a variable number of adenines. This suggests that the REase gene was excised from the genome. The same sequence was found in *H. cetorum* and *H. acinonychis*, the closest known relatives of *H. pylori* (Supplementary Table 3 and Supplementary Figure 1). Moreover, the phylogenetic trees of MTase and REase gene sequences in general were congruent with the global population structure of *H. pylori* (Figure 1) (23). This implies that the R-M system was acquired early in the history of this gastric pathogen. The REase gene appears to have been lost later during species evolution in the majority of the strains, likely before the first modern humans left Africa. Nonetheless, the REase gene could have been reintroduced in some strains (i. e. hpEurope strains) via recombination of the flanking repeats.

**Figure 1.**
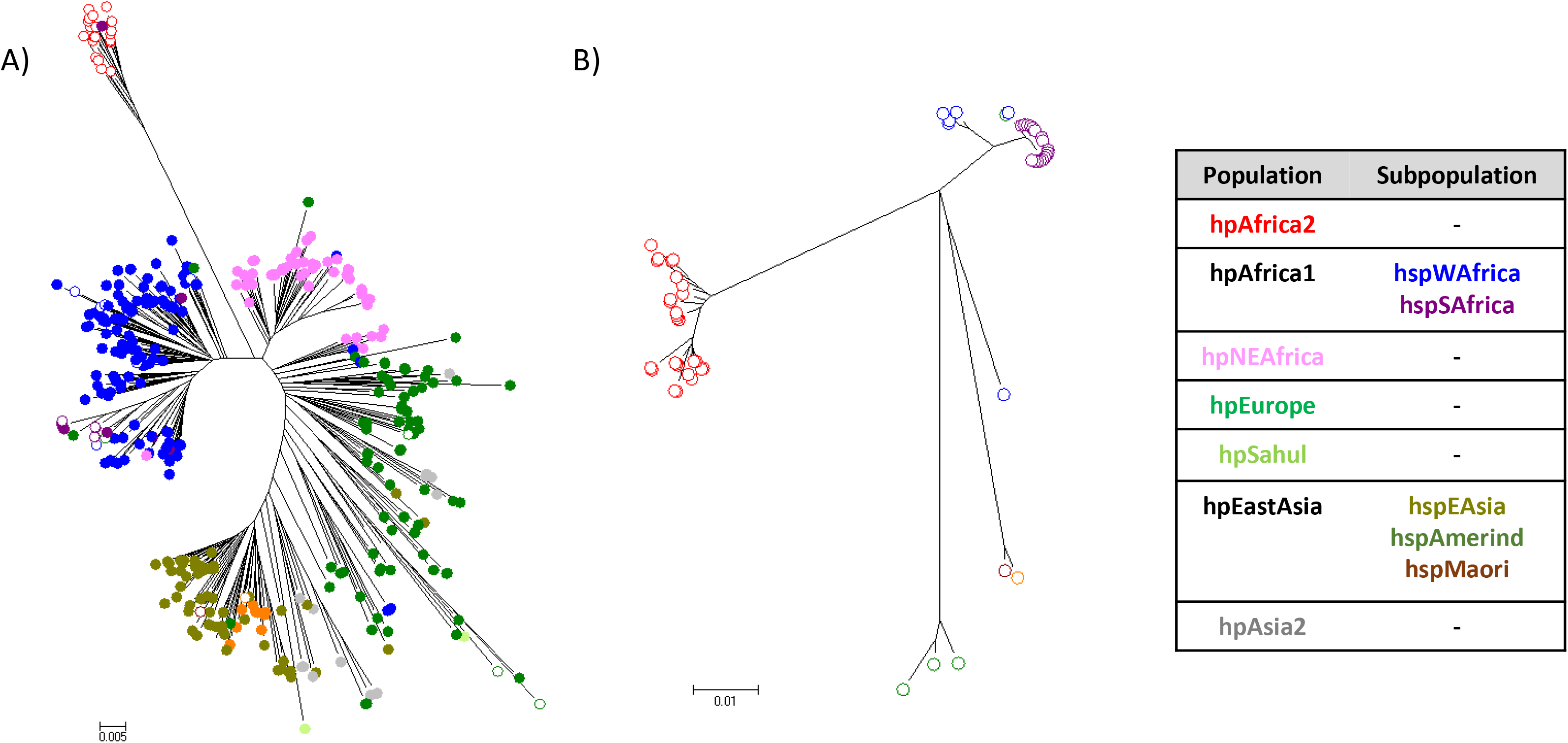
Phylogenetic analysis of the GCGC-specific R-M system JHP1050/1049 (M. Hpy99III/Hpy99III) in *H. pylori*. Neighbour-Joining trees based on the MTase M. Hpy99111 (A) and the REase Hpy99III (B) nucleotide sequences. In both cases, strain symbols are coloured according to the phylogeographic population assignment based on seven gene MLST and STRUCTURE analysis. Circles represents strains without REase gene, while diamonds are used for strains containing both MTase and REase genes.

### Construction of MTase mutants and analysis of target motif abundance

To functionally characterize this highly conserved MTase, we constructed MTase-deficient mutants. The MTase gene was disrupted in the strains 26695 (hpEurope), H1 (hspEAsia) and BCM-300 (hspWAfrica) and the whole R-M system was inactivated in strain J99 (hspWAfrica), the only of the four strains that contained both MTase and REase. Genes were inactivated by insertion of an antibiotic resistance cassette. The loss of methylation was verified by restriction assays using the restriction enzyme HhaI that can only cleave unmethylated GCGC sequences (Supplementary Figure 2). In the following text, mutants are named by the wild type strain name followed by –mut. Complementation of the MTase in 26695 and in J99 was performed by reintroducing the MTase gene of 26695 (see Material and Methods). The transcription of the MTase gene was tested in the four strains, and found to vary substantially between strains (Supplementary Figure 3), whether these differences between mRNA amounts have any functional implications is currently unknown.

Methylome comparison of the 4 strains exhibited only 4 methylated motifs shared between the strains (G^m5^CGC, G^m6^ATC, C^m6^ATG and G^m6^AGG) (Supplementary Table 4). The G^m5^CGC motif was very common in all four genomes, although the number of motifs differed between strains (Table 1). The distribution of motifs among the genomes was not uniform. We compared this observed distribution to the motif density that would be expected from a random distribution of motifs across the genomes. While the number of motifs was generally higher than expected for a random distribution, fewer motifs than predicted were found in the cagPAI and the plasticity zones (PZ) (Supplementary Figure 4A). Finally, we calculated the total number of GCGC motifs that would randomly occur in the genomes, the coding regions and the intergenic regions according to the nucleotide composition of *H. pylori*. The observed number of motifs in the whole genome and in the coding regions was higher than the expected number of motifs, while the calculated motifs in intergenic regions were similar to the observed number (Table 1). Therefore, coding sequences appear to display an over-representation of motifs.

**Table 1.** Observed and expected frequencies of GCGC motifs in the genome sequences of the four *H. pylori* strains analysed in this study.

| Strain | Genome size (bp) | Total length of CDS (bp) | Total length of intergenic sequences (bp) | Predicted no. of GCGC sites/1kb | No. of motifs in genome | Expected no. of motifs in genome | No. of motifs in CDS | Expected no. of motifs in CDS | No. of motifs in intergenic sequences | Expected no. of motifs in intergenic sequences |
| --- | --- | --- | --- | --- | --- | --- | --- | --- | --- | --- |
| 26695 | 1667867 | 1494807 | 173060 | 3.76 | 6269 | 2669 | 5950 | 2392 | 319 | 277 |
| J99 | 1643831 | 1486413 | 157418 | 3.89 | 6399 | 2630 | 6110 | 2378 | 289 | 252 |
| H1 | 1563305 | 1436409 | 126896 | 3.74 | 5846 | 2501 | 5655 | 2298 | 191 | 203 |
| BCM-300 | 1667883 | 1520688 | 147195 | 3.91 | 6523 | 2669 | 6273 | 2433 | 250 | 236 |

**Figure 4.**
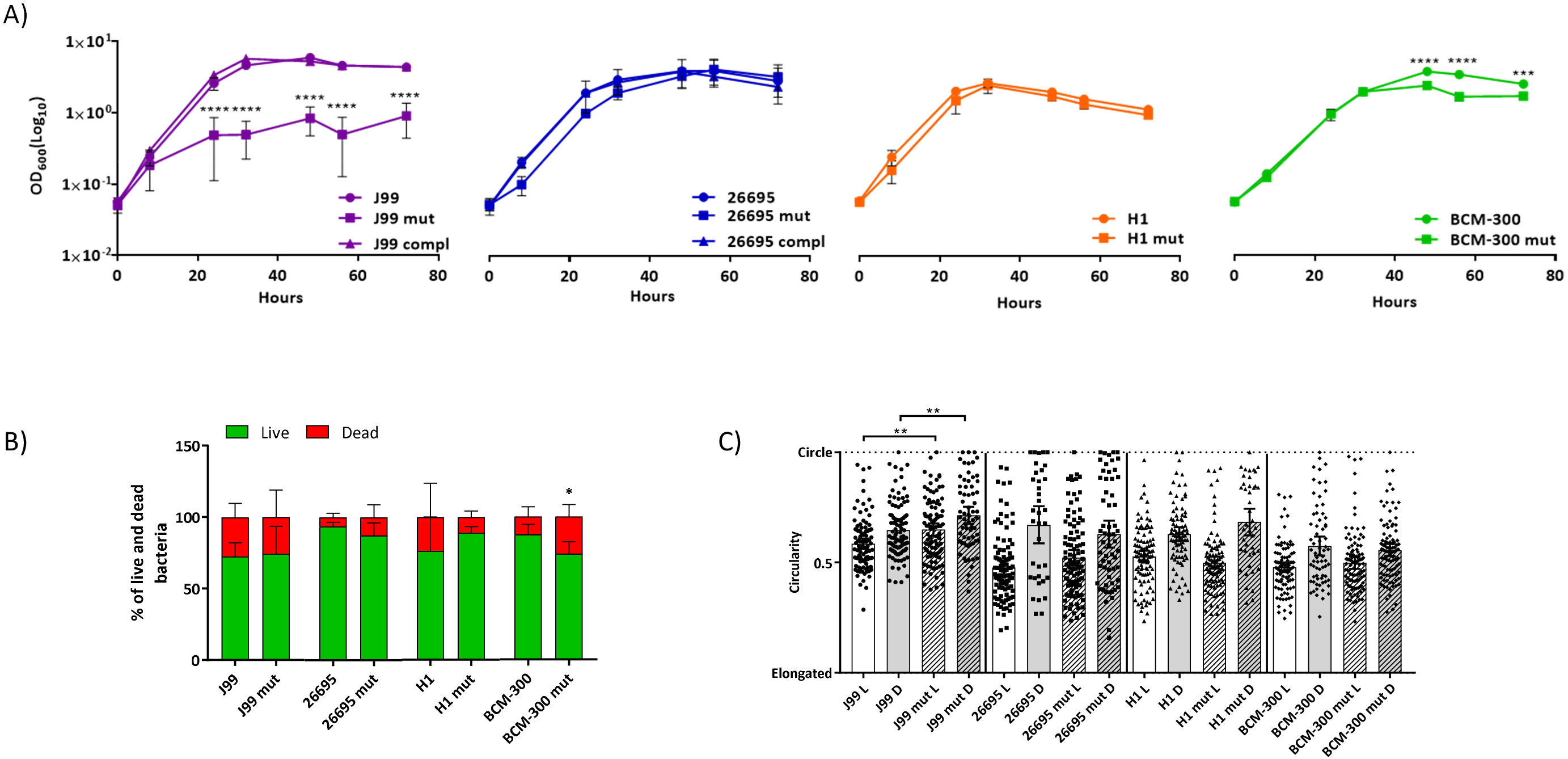
MTase JHP1050 inactivation causes phenotypic effects that vary between strains: Growth, viability and morphology. A) Growth curves for four wild type strains and mutants were measured until 72 hours. A significant growth defect was observed for J99-mut when compared with J99 wt and the complemented strains. Statistics: 2-way ANOVA, p<0.05, bars: Standard-Deviation (SD). B) Viability of the strains was studied using epifluorescence microscopy after Live/Dead staining. Similar viability was observed in all the cases. Statistics: 2-way ANOVA, p<0.05, bars: SD. C) Bacterial morphology was quantitated using Image J from pictures of the epifluorescence microscopy. A value of 0 represents complete elongated bacteria, while a value of 1 means complete circle. In general, live bacteria (L) were more elongated than dead bacteria (D). J99-mut was significantly more rounded than J99 wt. Statistics: One-Way ANOVA, p < 0.05, bars: 95% Coefficient-Interval (CI)

### Comparative RNA-Seq transcriptome analysis of *H. pylori* J99 and BCM-300 and their isogenic MTase mutants

Due to the extraordinary conservation of the G^m5^CGC MTase in all analyzed strains despite the absence of a cognate REase, we postulated that the function of the enzyme might be more important than simply serving for self-DNA protection. Therefore, in order to study a putative role in gene regulation, we performed comprehensive RNA-Seq analysis in the strains J99, BCM-300 and the two corresponding isogenic MTase mutants.

Whole transcriptome comparison of the J99-mut and J99 wt strains exhibited 225 differentially expressed genes (DEGs).115 genes were upregulated and 110 downregulated in J99-mut compared with J99 wt (p-adjusted value < 0.01, Fold Change (FC) > 2). In contrast to J99, the transcriptomes of the BCM-300-mut and wt strains showed only 29 genes that were differentially expressed in the mutant, all of which were downregulated (p-adjusted value < 0.01, FC > 2) (Supplementary Table 5). The two mutants, J99-mut and BCM-300-mut, shared 10 downregulated genes but no upregulated genes (Table 2). Using qPCR, we confirmed some of the shared genes were significantly downregulated as shown by RNA-Seq (Supplementary Figure 5E, 5F).

**Table 2.** Shared differentially expressed genes (DEG), displaying GCGC methylation-dependent transcription in *H. pylori* J99 and BCM-300. Positive values for Fold Change (FC) indicate lower transcription in the mutants compared to the wt strains.

| Gene | Description | J99 locus_tag | J99 FC | BCM-300 locus_tag | BCM-300 FC |
| --- | --- | --- | --- | --- | --- |
| *bioD* | Dethiobiotin synthetase | *jhp_0025* | 2.1986 | *BCM_00034* | 2.9978 |
|  |  |  |  | *BCM_00035* | 2.9424 |
| *feoB* | iron(II) transport protein | *jhp_0627* | 3.8803 | *BCM_00707* | 4.3250 |
| - | unknown | *jhp_0749* | 3.8245 | *BCM_00859* | 3.1947 |
| *moeB* | molybdopterin/thiamine biosynthesis activator | *jhp_0750* | 4.0863 | *BCM_00860* | 3.6033 |
| - | unknown | *jhp_1102* | 2.4868 | *BCM_01112* | 2.2810 |
| *cah* | Alpha-carbonic anhydrase | *jhp_1112* | 2.0723 | *BCM_01124* | 3.3563 |
| *trmU* | tRNA-methyltransferase | *jhp_1254* | 4.5288 | *BCM_01276* | 5.7005 |
| - | unknown | *jhp_1281* | 3.4690 | *BCM_01305* | 2.0216 |
| - | unknown | *jhp_1253* | 2.9141 | *BCM_01275* | 3.2789 |
| *crdR* | Response regulator | *jhp_1283* | 2.8855 | *BCM_01307* | 3.2789 |
|  |  | *jhp_1443* | 2.9141 |  |  |

In order to understand how the distribution of motifs could play a role in transcriptional regulation, we analysed 500 bp sequence upstream of each DEG in comparison with sequences upstream of genes that were not differentially regulated (non-DEGs), and with the number of motifs within CDS.

In strain BCM-300, the number of GCGC motifs located within 500 bp upstream of the start codon was higher for the 29 DEGs than for the genes that were not differentially regulated (Figure 2A). In contrast, in strain J99, the percentage of genes with three or more GCGC motifs within 500 bp upstream of the start codon was similar for DEGs and non-DEGs (Figure 2C). However, the 10 DEGs of strain J99 that were shared with BCM-300 showed the same overrepresentation of GCGC motifs observed in strain BCM-300 (Figure 2B, 2D). Furthermore, DEGs in BCM-300 displayed more motifs within their CDS than expected if GCGC motifs were distributed randomly across the whole genome, while the opposite effect occurred for the non-DEGs. The same trend was evident in J99 when we only compared the DEGs shared with BCM-300 with the rest of the genes (Supplementary Figure 6A).

**Figure 2.**
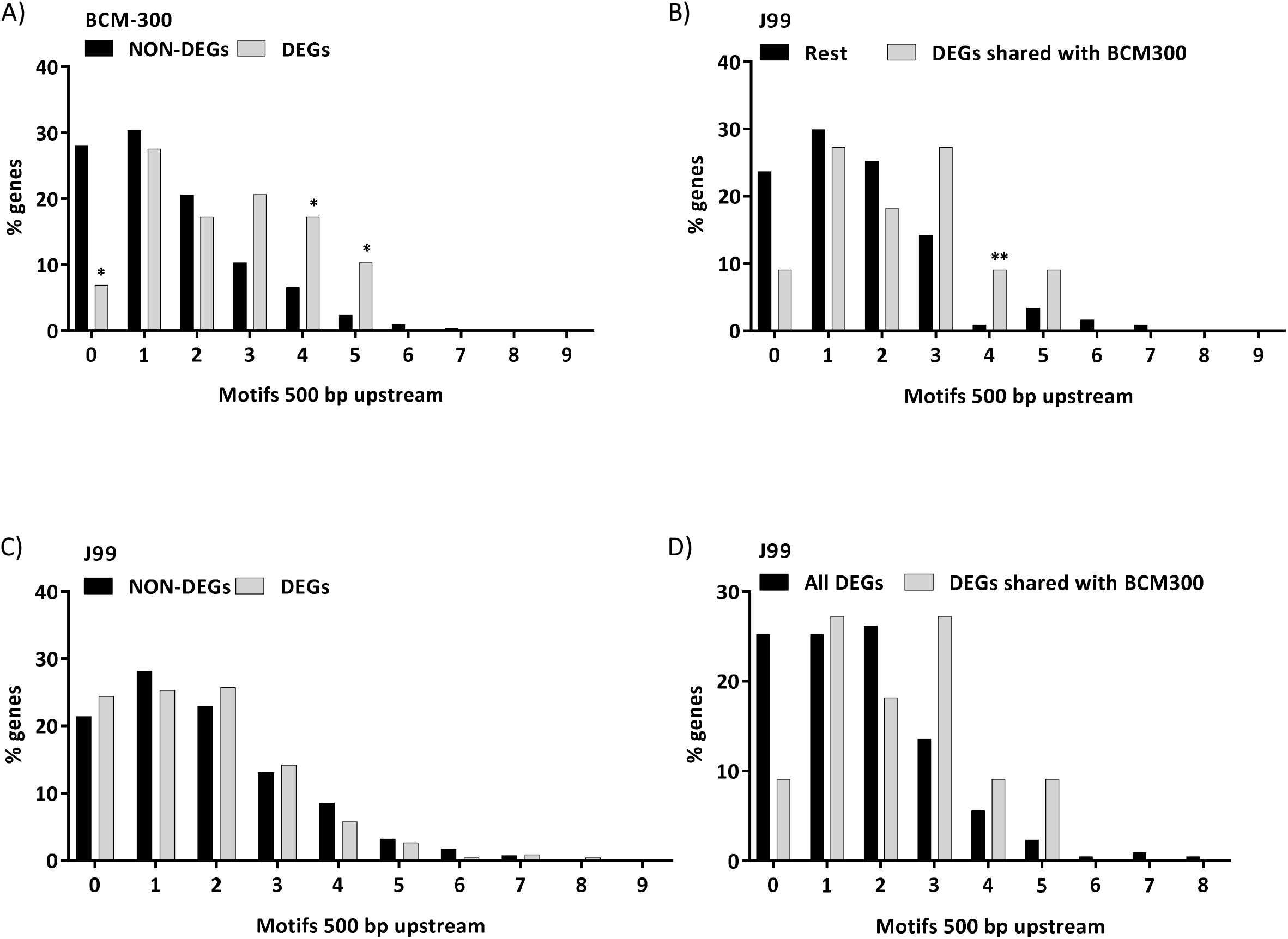
Graphical representation of the percentage of genes with GCGC motifs 500 bp upstream of the start codon. A) Percentage of Non-DEGs vs DEGs with motifs 500 bp upstream in BCM-300. B) DEGs in J99 shared with BCM-300 vs the rest of the genes in J99. C) Non-DEGs vs DEGs with motifs 500 bp upstream in J99. D) DEG in J99 shared with BCM-300 vs the rest of the DEG genes in J99 (All DEG). Statistics: Chi-square, p < 0.05.

In addition, we observed that of the 10 shared DEGs, 6 harboured GCGC motifs within the 50 bp upstream of the TSS described by Sharma and colleagues in strain 26695 (51), called here upstream region of the TSS (upTSS).

Sequences within the putative promoter regions immediately upstream of the TSS are likely to exert the strongest influence on transcriptional regulation. We compared the upTSS of 26695 with J99 and BCM-300 via sequence alignment. There were 48 genes in J99 and 45 in BCM-300 with motifs within the 50 bp upstream sequence (sRNA and asRNA were excluded). In J99, of the 225 DEGs, 13 genes harboured GCGC motifs within the upTSS sequence. In BCM-300, 11 of the 29 DEGs carried motifs within the upTSS. This proportion of DEGs with motifs within the upTSS suggests that the window of 50 bp upstream of the TSS may play a role in transcription regulation. Indeed, the FC was slightly increased by motifs within the upTSS (Supplementary Figure 6B).

### Direct regulation of gene expression by ^m5^C methylation

Inactivation of the M. Hpy99111 MTase had different effects on the transcriptomes of the two strains tested, with far more genes affected in strain J99 vs. the BCM-300 strain. We hypothesized that the loss of GCGC methylation might have both direct and indirect effects on transcription. In order to demonstrate a direct association between methylation and gene expression, we generated a set of mutants of strain J99 where site-specific mutations were introduced into selected GCGC motifs located within the CDS as well as in the upstream region of one gene showing strong differential regulation.

The selected gene for this approach *(jhp0832*) was downregulated in J99-mut (FC = 5.95). Its homolog in *H. pylori* strain 26695 was reported to be an antitoxin from a Type II Toxin-Antitoxin (TA) system (52). The cognate toxin *(jhp0831*) was also downregulated in J99-mut (FC = 3.64). The two genes belong to the same operon where the antitoxin is located upstream of the toxin. No homologous genes were found in BCM-300.

Two GCGC motifs were located within the 500 bp upstream window of the antitoxin gene and one within the coding sequence. Of the two motifs upstream, one was located within the upTSS in J99 and overlapped the -10 box of the predicted promoter (Figure 3). Thus, due to the high FC and the distribution of motifs upstream and within the gene sequence, *jhp0832* seemed to be a good candidate to test the GCGC motif-dependent regulation.

**Figure 3.**
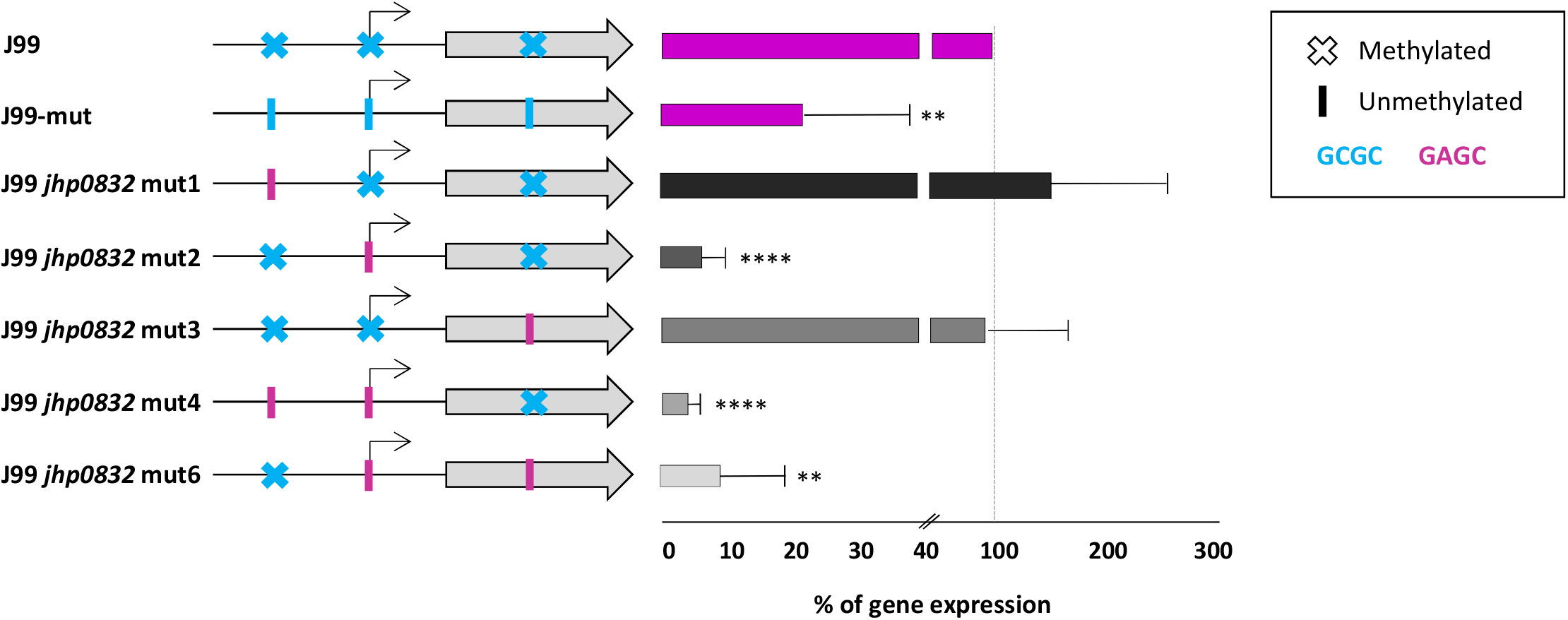
Quantification of the transcription of *jhp0832* in *H. pylori* strains J99, J99-mut and the J99 mutants with point mutations within the GCGC motifs. qPCR results are represented in the right panel, 3 different biological replicates were performed. J99-mut and the 3 mutants with the GCGC motif mutated within the promoter sequence had a significantly lower expression of the gene compared to J99 wt in all the replicates. Instead, the other two mutants displayed an altered expression that did not follow a regular pattern, since the expression differed among replicates. Statistics: One-Way ANOVA, p < 0.05, bars: SD. Legend: The gene is shown as a gray arrow. The predicted promoter is represented by a black arrow. Crosses represent methylated motifs while vertical lines mean unmethylated motifs. The GCGC motifs appear in blue and the mutated motifs to GAGC are colored in pink.

We constructed three mutants where each of the motifs was individually changed to GAGC so that the motif could no longer be methylated *(jhp0832* mut1, *jhp0832* mut2 and *jhp0832* mut3). We also constructed two mutants *(jhp0832* mut4 and *jhp0832* mut6) where 2 out of the 3 GCGC motifs were mutated (Figure 3, Table 3). We were unable to generate a mutant carrying mutations in all 3 motifs.

**Table 3.** List of mutants carrying different point mutations modifying the GCGC motifs within or immediately upstream of *jhp0832*. All mutants were constructed using the MuGent technique (see Materials and Methods) using the indicated plasmids and the rdxA∷CAT PCR product.

| Mutant name | GCGC motif mutated | Plasmid | Antibiotic resistance cassette |
| --- | --- | --- | --- |
| *jhp0832* mut1 | 1 | pSUS3427 | CAT |
| *jhp0832* mut2 | 2 | pSUS3428 | CAT |
| *jhp0832* mut3 | 3 | pSUS3429 | CAT |
| *jhp0832* mut4 | 1,2 | pSUS3427, pSUS3428 | CAT |
| *jhp0832* mut6 | 2,3 | pSUS3428, pSUS3429 | CAT |

Differential expression of *jhp0832* was determined by quantitative PCR (qPCR). Three of the mutants (*jhp0832* mut2, *jhp0832* mut4 and *jhp0832* mut6) displayed a strong downregulation of *jhp0832* expression, similar to J99-mut. Interestingly, these mutants shared the mutation in the G^m5^CGC motif located within the upTSS and the predicted promoter of the gene. In contrast, modification of the motifs outside of the upTSS did not consistently alter the expression of the gene (Figure 3).

### Phenotypes of *H. pylori* GCGC MTase mutants: growth, viability and shape

In order to test whether the absence of ^m5^C methylation and the associated differential transcriptomes had a role in the fitness of *H. pylori*, we determined the growth of the strains in liquid medium (Figure 4A). J99-mut had a significant growth defect compared with the J99 wild type strain.

Complementation of the MTase gene restored the observed growth phenotype. Similarly, a significant reduction in growth was shown for BCM-300-mut at stationary phase. Although non-significant, a slight delay in growth was noted in 26695-mut and H1-mut compared to the wild type and the complemented strains.

Bacterial morphology serves to optimize biological functions and confers advantages to particular niches.*H. pylori* is a spiral-shaped bacterium that can enter a coccoid state under certain stress conditions (53).*H. pylori* J99-mut entered a coccoid state very early in liquid cultures. A substantial proportion of coccoid forms were visible between 6-9 hours after inoculation while they are rarely found in the wild type strain at this time point (Supplementary Figure 7A). An effect of the inactivation of JHP1050 on the morphology was not observed for the other three strains 24 hours post-inoculation (Supplementary Figure 7B). Complementation of J99-mut restored the wild type phenotype. We note that Live/Dead staining did not show a significant difference between the percentage of live vs. dead bacteria between the wild type and the mutant strains collected from 22-24 hour plates. There was a slight reduction in viability in the BCM-300-mut strain, but no differences were found in the other strains (Figure 4B). As in the liquid cultures, an increased number of rounded bacteria were noticed for J99-mut (Figure 4C).

### ^m5^C methylation contributes to the high mutation frequency in *H. pylori*

*H. pylori* lacks most of the genes involved in mismatch repair (MMR) in other bacteria which is thought to be at least partially responsible for the high mutation rate of this bacterium (54,55). Deamination of ^m5^C to thymine (T) is responsible for the most common single nucleotide mutation (56).*H. pylori* is known to have a very high mutation rate, and ^m5^C MTases might contribute to that by increasing the number of nucleotides susceptible to deamination. To test whether ^m5^C methylation within GCGC motifs played a role in *H. pylori* evolution by favouring deamination, we aligned whole genomes of two *H. pylori* strains (26695 and PeCan18), used as reference, against 11 other complete genome sequences (see Material and Methods for details). The results strongly supported a role of ^m5^C methylation in *H. pylori* mutagenesis, since the percentage of C->T mutations within G^m5^CGC motifs was significantly higher than the overall C->T or C-> another base transition in the genomes of all the tested strains. Therefore, the ^m5^C methylation of the common GCGC motif in all *H. pylori* strains may contribute to the high mutation rate of *H. pylori* and its overall low GC content by favouring deamination (Supplementary Figure 4B).

### Regulation of Outer Membrane Proteins (OMPs) and adherence by G^m5^CGC methylation is strain-specific

OMP genes represent approximately 4% of the *H. pylori* genome (57). Fourteen OMPs were found to be upregulated in J99-mut (Supplementary Table 5). Only three of these OMPs were slightly upregulated in BCM-300-mut but the FC was lower than the cut-off of 2. Confirmation of the upregulation of OMP genes was performed using qPCR in J99-mut (Supplementary Figure 5C, D). We detected either no regulation or weak upregulation in the other three mutated strains (Supplementary Figure 5C, D), which was in agreement with the transcriptome data obtained for BCM-300. A bacterial adherence assay based on coincubation of fixed AGS cells with all four wild type strains and corresponding isogenic GCGC mutants was performed. Only J99-mut had a significantly higher adherence to the cells compared to the respective wild type strain, while no significant differences in adherence were determined for the rest of the strains (Figure 5C). Taken together, the increased expression of a number of OMP genes in the absence of methylation in J99 might contribute to a stronger adherence of the bacteria to the cells.

**Figure 5.**
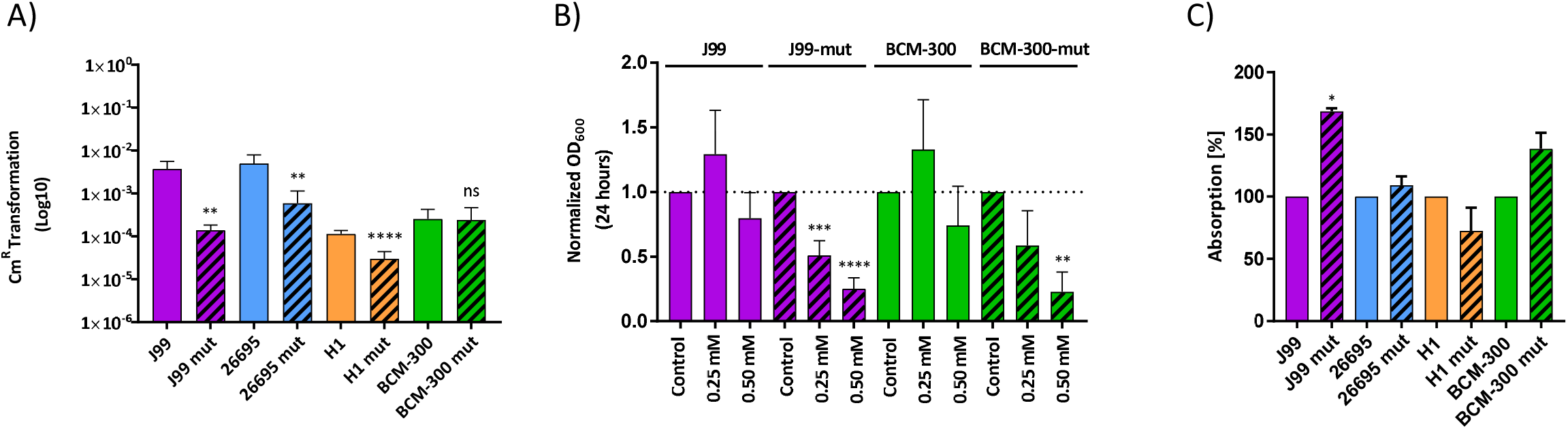
MTase JHP1050 inactivation causes phenotypic effects that vary between strains: Natural competence, resistance to copper, and adherence to host cells. A) Transformation experiments were performed transforming bacteria with 1 μg/ml of gDNA. Lower transformation frequencies were observed for three of the mutated strains. No significant difference was observed for BCM-300 Statistics: Welch’s unpaired t test, p < 0.05, bars: SD. B) The growth of J99 wt, J99-mut, BCM-300 wt and BCM-300-mut strains was measured 24 hours post-inoculation after addition of different concentrations of copper sulphate to the cultures. A clear growth defect can be accounted for the mutated strains when there is an excess of Cu. Data was normalized to a control culture without copper. Statistics: One-Way ANOVA, p<0.05, bars: SD. C) Adherence of *H. pylori* wt and mutant strains to fixed AGS cells. An increase in cell adherence was observed for J99-mut, corresponding to the upregulation of multiple OMP genes in the MTase mutants. Statistics: unpaired t test, p < 0.05, bars: SD

### GCGC methylation regulates natural competence in *H. pylori*

Natural competence is a hallmark of *H. pylori*. Competence is conferred by the ComB system, an unusual type IV secretion system related to the VirB system of *Agrobacterium tumefaciens* (58). RNA-Seq results identified three *com* genes *(comB8, comB9* and *comEC)* that were less transcribed in J99-mut compared to the wild type strain, but the genes were not found to be differentially regulated in BCM-300. ComB9 and ComB8 are part of the outer- and inner-membrane channels of the DNA uptake system, while ComEC allows the translocation of the DNA through the inner membrane to the cytoplasm. qPCR confirmed the downregulation of these genes in 26695-mut and H1-mut strains in comparison with the respective wild type strains (Supplementary Figure 5A, 5B).

The DNA uptake capacity of the four mutated strains was quantitated by counting recombinant colonies carrying an antibiotic resistance cassette after standardized transformation experiments (see Materials and Methods). A significant reduction in the efficiency of transformation to chloramphenicol resistance was observed in the J99, 26695 and H1 mutants compared to their respective wild type strains, but no difference was apparent for BCM-300 (Figure 5A). The down-regulation of these three components of the ComB system might be sufficient to reduce the competence in three of the strains.

### Loss of ^m5^C methylation increases susceptibility to copper toxicity

Copper (Cu) is an essential metal used by *H. pylori* as a cofactor in multiple processes and it has been shown, for example, to be important for colonization (59). However, an excess of heavy metals can be toxic for the bacterial cells, leading to the existence of several mechanisms to control Cu homeostasis. One of the mechanisms involves the two-component system CrdR/S. In the presence of Cu, the sensor kinase CrdS phosphorylates the response regulator CrdR triggering the activation of a copper resistance protein and a copper efflux complex (60).

The transcriptional regulator gene *crdR* was less expressed in both J99 and BCM-300 MTase mutants (Table 2). In both strains, one GCGC motif is located within the upTSS of the transcriptional regulator, suggesting a direct regulation via ^m5^C methylation. To test whether the mutated strains were less resistant to Cu due to the lower expression of the *crdR* gene, we compared the influence of added copper sulphate on growth in liquid culture between MTase mutants and wild type strains. The presence of Cu caused a clear growth defect of the mutants when compared with the wild type strains, and with a control culture without added Cu (Figure 5B). The results indicates that ^m5^C methylation within the upTSS is required to ensure sufficient transcription of the transcriptional regulator to protect against an excess of copper.

## DISCUSSION

Most previous studies of R-M systems in *Helicobacter pylori* have focussed on the striking diversity of methylation patterns and its implications. In contrast to the dozens of MTases only present in subsets of strains, *H. pylori* also possesses few enzymes that are highly conserved between strains. Here, we have explored the function of one ^m5^C MTase (JHP1050) that we predicted to be active in all of a globally representative collection of 459 *H. pylori* strains analysed. The collection included isolates from the most ancestral *H. pylori* population, hpAfrica2, and the presence of the MTase in all *H. pylori* phylogeographic populations and subpopulations indicates that the gene has been part of the *H. pylori* core genome since before the Out of Africa migrations, and before the *cag* pathogenicity island was acquired (23). The cognate REase gene was detected in few strains only, almost all of which belong to African *H. pylori* populations. This indicates that the REase was excised from the genome very early in the history of this gastric pathogen. These data designate strong selective pressure to maintain the activity of the MTase, while the REase gene either lost its function or was completely deleted. The apparent strong selection of the maintenance of this MTase in the *H. pylori* genome was in striking contrast to the cognate REase and to the vast majority of R-M systems so far identified in *H. pylori*, indicating that the MTase alone is likely to serve an important function for the bacterium. Since methylation has been shown to influence gene expression in several bacterial species, we considered a regulatory function most likely, and performed global transcriptome analysis using RNA-Seq.

The results obtained by RNA-Seq analysis of two *H. pylori* wild type strains, J99 and BCM-300, and their respective MTase mutants confirmed our hypothesis that GCGC methylation affects the transcription of multiple *H. pylori* genes, but we were surprised by the substantial differences between the two strains. While there were 225 DEGs in J99, whose transcription was significantly changed in the MTase mutant, only 29 genes showed an altered expression in BCM-300, and only 10 DEGs were shared between both strains.

To better understand the relationship between GCGC methylation and transcriptional gene regulation, we studied the correlation between the presence of GCGC motifs within coding sequences and upstream regulatory sequences and the effect of a loss of methylation on transcription.

DEGs were more likely to contain more than three motifs in the 500 bp sequence upstream of the start codon than the genes not showing significant differential regulation (Figure 2). Among the DEGs, the presence of GCGC motifs within the upTSS was significantly associated with higher fold change (FC) values (Supplementary Figure 6B). Moreover, there were more DEGs with higher number of motifs within the coding sequence than expected when compared with the non-DEGs (Supplementary Figure 6A). These results are similar to reports from *Vibrio cholerae*, where a significant correlation between differential regulation and the number of motifs within the coding sequence was reported for a ^m5^C MTase (21).

Six of the 10 DEGs shared between J99 and BCM-300 contained GCGC motifs within the upTSS. We therefore investigated the relationship between the presence of a methylatable GCGC sequence and gene transcription using site directed mutagenesis. When the methylated G^m5^CGC motif within the promoter of the DEG *jhp0832* was changed to a non-methylated GAGC motif, this caused a clear down-regulation of the transcription that was similar to the effect of MTase inactivation (Figure 3). This provides strong and direct evidence that methylation of the GCGC motif within a promoter sequence affects gene transcription. Similar findings were previously reported for G^m6^ACC motifs methylated by the *H. pylori* ModH5 MTase, which are involved in the control of the activity of the *flaA* promoter in strain P12 (39). The exact mechanism(s) how methylated sequence motifs within promoters and most likely also within coding sequences influence gene expression in *H. pylori* is still unknown. One emerging paradigm is exemplified by the essential cell cycle regulator GcrA from *Caulobacter crescentus*, a σ70 cofactor that binds to almost all σ70 promoters, but only induces transcription of genes that harbour G^m6^ANTC methylated sites in their promoters (61).

The 10 DEGs shared by both strains were less expressed in the absence of methylation. Thus, in contrast to eukaryotes, where CpG methylation in promoter regions leads to the silencing of genes, methylation of GCGC sites in *H. pylori* promoters enhances transcription. Many of the shared DEG belong to conserved cellular pathways (i. e. biotin synthesis, Fe(ii) uptake, molybdopterin biosynthesis, bicarbonate and proton production, tRNA modification) and also include a transcriptional regulator involved in copper resistance. Based on these observations, we propose that the conserved GCGC-specific MTase directly controls the expression of those genes involved in various, partially fundamental, cellular pathways.

The inactivation of the MTase caused a substantial growth defect and accelerated conversion to coccoid cells in *H. pylori* J99 that were restored to wild type growth in a complemented strain. The three other wild type strains investigated did not show a similar growth defect when the MTase was inactivated. Other phenotypic effects induced by the MTase inactivation were observed in all or multiple strains. They included functions important for virulence, such as morphology, competence and adherence to gastric epithelial cells. The genome diversity of *H. pylori*, the distribution of motifs among the genomes and the variable methylomes due to the activity of other MTases must influence global gene expression. It was demonstrated recently that deletions of two strain-specific MTases, the ^m5^C MTase M. HpyAVIB (62) and the ^m4^C MTase M2. HpyAII (63) both also had regulatory effects on the *H. pylori* transcriptome. While the effects differed widely from those observed for the M. Hpy99III MTase studied here, there were some genes differentially regulated by more than one MTase, suggesting that the effects of different MTases may be interlinked. Thus, the strain-specific phenotypes observed in the absence of ^m5^C methylation in GCGC motifs are likely to reflect the complex and intrinsic diversity of *H. pylori* at the genome, methylome, and transcriptome levels.

While we clearly showed that methylation of a GCGC motif overlapping the promoter within the upTSS directly affected transcription, we currently do not understand how the presence or absence of GCGC methylation can affect so many genes in strain J99, and which mechanisms contribute to strain-variable effects. It seems likely that at least some of the massive changes observed in strain J99 are indirect effects, e. g. resulting from the downregulation of genes affecting growth. The effect of MTase inactivation in any given strain is likely to be the net outcome of interlinked direct and indirect regulatory effects that will need to be further elucidated in the future. Methylation may affect DNA topology, which has a strong influence on genome-wide gene regulation, causing secondary effects on the global transcriptome by a plethora of mechanisms. For example, modifications of DNA topology affect the binding of DnaA to the OriC2 of *H. pylori* (64). The *flaA* promoter, whose expression is governed mainly by the transcription factor σ^28^, was shown by extensive mutagenesis t be strongly modulated in a topology-dependent manner during the growth phase (65). This also fits to the previously described methylation-dependent indirect regulation of the *flaA* promoter (39). Finally, several direct and indirect means of methylation-mediated regulatory mechanisms might not exclude each other, generating an intricate network fine-tuning gene expression, which depends on genome-wide methylation.

## CONCLUSION

Global changes in ^m5^C DNA methylation patterns in *H. pylori* affect the expression of several genes directly or indirectly, which results in both strain-independent (conserved) and strain-dependent effects. Motifs situated within promoter sequences have a direct effect on transcription, while surrounding motifs might modulate the expression indirectly by, for example, altering the topology of the DNA. Furthermore, methylation of G^m5^CGC target sequences maintains regulated the transcription of genes involved in metabolic pathways, competence and adherence to gastric epithelial cells.

## ACCESSION NUMBERS

RNA-Seq data will be made publicly available prior to publication in a peer-reviewed journal.

## ACKNOWLEDGEMENT

We thank Sandra Nell for help with assembling the collection of 459 globally representative *H. pylori* genomes, and Gudrun Pfaffinger for excellent technical assistance.

